# Neuropilin 1 mediates epicardial activation and revascularization in the regenerating zebrafish heart

**DOI:** 10.1101/468504

**Authors:** Vanessa Lowe, Laura Wisniewski, Jacob Sayers, Paul Frankel, Nadia Mercader-Huber, Ian C Zachary, Caroline Pellet-Many

## Abstract

Unlike adult mammals, zebrafish are able to naturally regenerate their heart. A key mechanism in zebrafish heart regeneration is the activation of the epicardium, leading to the establishment of a supporting scaffold for newly formed cardiomyocytes, angiogenesis and cytokine secretion. Neuropilins (NRPs) are cell surface co-receptors mediating functional signaling of kinase receptors for cytokines known to play critical roles in zebrafish heart regeneration, including Platelet-Derived growth factor (PDGF), Vascular Endothelial growth factor (VEGF), and Fibroblast growth factor (FGF). Herein, we investigated the role of neuropilins in the response of the zebrafish heart to injury and its subsequent regeneration. All four zebrafish neuropilin isoforms, *nrp 1a*, *1b*, *2a*, and *2b*, were upregulated following cardiac cryoinjury and were strongly expressed by the activated epicardium. A *nrp1a* mutant, coding for a truncated, non-functional protein, showed a significant delay in heart regeneration in comparison to Wild-Type fish and displayed persistent collagen deposition. The regenerating hearts of *nrp1a* mutants were less vascularized and epicardial-derived cell migration and re-expression of the developmental gene Wilms’ tumor 1 was severely impaired in *nrp1a* mutants. Moreover, cryoinjury-induced activation and migration of epicardial cells in heart explants was strongly reduced in *nrp1a* mutant zebrafish. These results identify a key role for Nrp1 in zebrafish heart regeneration, mediated through epicardial activation, migration and revascularization.

## Introduction

Ischemic heart disease remains the leading cause of death worldwide and although improved therapeutic treatments have led to an increase in myocardial infarction (MI) survival rates (1-3), cardiac function often remains severely compromised because adult mammalian hearts replace damaged tissue with an irreversible fibrotic scar (4, 5). This often leads to the development of chronic heart failure, further MIs, and fatal arrhythmias (6). In contrast to mammals, zebrafish have the remarkable ability to regenerate lost or damaged cardiac tissue *via* cardiomyocyte proliferation and resorption of fibrotic tissue, ultimately restoring cardiac function (7-10). Understanding the underlying mechanisms that govern zebrafish heart regeneration may identify therapeutic targets important for stimulating cardiac repair following MI in mammals.

Zebrafish heart regeneration involves a well described but incompletely understood sequence of cellular processes and signaling events. The epicardium, a mesothelial cell monolayer encasing the heart, has been strongly implicated as a key regulator of the regenerative response (11-13). Upon cardiac damage, the epicardium is activated (14, 15), undergoing proliferation and secreting cytokines that stimulate cardiomyocyte cell cycle re-entry (16). Autocrine and paracrine signals induce a subpopulation of epicardial cells to undergo a process known as epithelial to mesenchymal transition (EMT) (17-21). These epicardial cells adopt an embryonic-like gene expression profile (22, 23), migrate into the injured region, and differentiate into fibroblasts and mural cells that support revascularization (17, 24). Some of the signaling pathways required for the epicardial regenerative response in zebrafish have been identified and characterized. In particular, platelet-derived growth factor (PDGF)-BB and fibroblast growth factor (FGF) are both essential for epicardial EMT and coronary neovascularization in the regenerating zebrafish heart (17, 18). Vascular endothelial growth factor (VEGF) was also found to play a key role in the early revascularization of the injured area (25).

PDGF, FGF, and VEGF are all ligands for neuropilin (NRP) transmembrane receptors (26-28). NRP1 and NRP2 share similar homology domain organization, with a large extracellular region essential for ligand binding, a single transmembrane domain, and a short cytoplasmic domain (29). NRP1 was first identified as a regulator of angiogenesis and neurogenesis mediated *via* VEGF and semaphorin3A, respectively (30-35). In zebrafish, it is also required for vascular development and is a mediator of VEGF-dependent angiogenesis (36). Furthermore, NRPs have been shown to mediate signaling pathways for other cytokines, including PDGF, FGF, and TGF-β in various tissues in both physiological and pathological settings (37-41). NRPs have also been reported to play a role in EMT in carcinomas (39, 42, 43), but, despite their known interactions with cytokines implicated in EMT, their role in the epicardial response to cardiac injury is currently unknown. We used the zebrafish heart cryoinjury model (7) to investigate the spatio-temporal expression of the four zebrafish neuropilin isoforms (*nrp1a*, *nrp1b*, *nrp2a*, and *nrp2b*, orthologues of human NRP1 and NRP2, respectively) in the regenerating heart. We show that all are upregulated in response to cryoinjury, with distinctive endocardial and epicardial expression during the regenerative response. Neuropilins are expressed in activated epicardial cells and zebrafish expressing a truncated loss-of-function Nrp1a (*nrp1a^sa1485^*) show impaired epicardial response to injury and exhibit reduced epicardial cell migration. Moreover, the revascularization of the injured area is also impaired in mutant fish. These findings reveal an essential role for Nrps in zebrafish heart regeneration, mediated by a new function for Nrps in epicardial activation and cell movement.

## Results

### Neuropilins are up-regulated during zebrafish heart regeneration

We quantified *nrp1a*, *nrp1b*, *nrp2a*, and *nrp2b* mRNA levels in whole ventricles following cardiac cryoinjury by absolute RT-qPCR and compared their expression with that in sham-operated hearts. *Nrp1a*, *nrp1b*, and *nrp2a* genes were up-regulated in injured hearts compared to sham-operated hearts early in the regenerative process (1 and 3 days post-cryoinjury) and returned to endogenous basal levels thereafter (Fig. 1A). In line with previous publications (44), we observed that *nrp2b* is the most highly expressed isoform in the heart under control conditions (Fig. 1A). However, qPCR did not show any significant *nrp2b* changes following cardiac damage, probably because any localized or cell type-specific cardiac up-regulation of this isoform was masked by its high basal expression.

**Figure 1.**
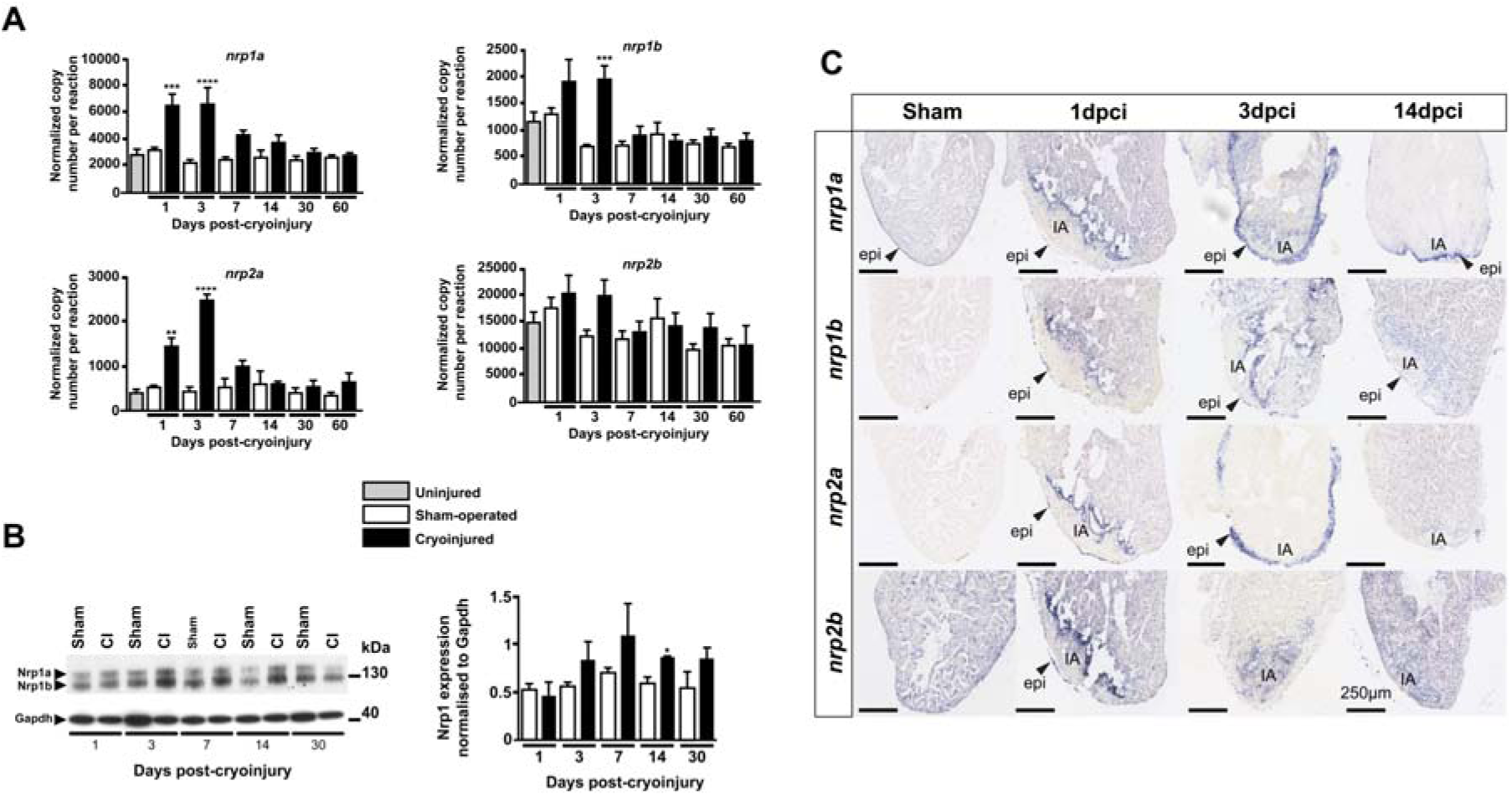
Neuropilins are upregulated during zebrafish heart regeneration. **(A)** Absolute quantitative PCR analysis of neuropilins at 1, 3, 7, 14, 30, and 60 days following cryoinjury or sham surgery. Basal expression was evaluated in uninjured hearts of age-matched wild type fish. Bars represent normalised copy number per reaction means ± S.E.M, ^⋆⋆^*p*<0.01, ^⋆⋆⋆^*p*<0.005, ^⋆⋆⋆⋆^*p*<0.001 (n = 4-5 with each *n* being a pool of 5 ventricles). **(B, left)** Adult zebrafish ventricle lysates 1, 3, 7, 14 and 30 days following surgery immunoblotted for Nrp1 and Gapdh. **(B, right)** Western blot quantification of Nrp1 protein in sham and cryoinjured ventricles 1, 3, 7, 14 and 30 days following surgery (n = 4-5, with each *n* being a pool of 3 ventricles). **(C)** *In situ* hybridization with digoxigenin-labelled anti-sense riboprobes were used to detect *nrp* isoforms in sham-operated and cryoinjured adult zebrafish hearts 1, 3 and 14 days post-cryoinjury (dpci). IA - injured area, epi-epicardium. Arrows indicate gene expression within the epicardium. Scale bar 250μm (*n* ≥ 3).

We also analyzed the expression of molecules implicated in Nrp-mediated signaling pathways and others with a known role in zebrafish heart regeneration (Fig.S1). Consistent with previous work, *pdgfrfβ* (NM_001190933), *pdgf*-*ab* (NM_001076757), and *tgfβ1a* (NM_212965) were all upregulated early after cryoinjury (Fig.S1) (10, 17). Because Neuropilins are VEGF co-receptors, we examined the regulation of *vegfaa* (NM_001190933), *vegfc* (NM_205734), and the VEGF receptors *kdrl* (NM_131472) and *flt1* (NM_001014829 and NM_001257153). In accordance with previous published data (45), *vegfc* was significantly upregulated following cardiac cryoinjury (Fig.S1), but we could not detect any significant change in the expression of *vegfaa*, *kdrl*, and *flt1.* In this context and at this time following the injury, *vegfc* is likely to be involved in inflammation and lymphangiogenesis as previously reported (46).

Nrp1 protein expression in zebrafish ventricles, detected by Western blot, was observed as two bands of approximately 130 kDa and 150 kDa, corresponding to Nrp1a (NM_001040326, 916 amino acids) and Nrp1b (AY49341, 959 amino acids), respectively (Fig. 1B). From 3 days post cryoinjury (dpci) and later, immunoblotting revealed an up-regulation of Nrp1 proteins in the injured hearts compared to sham-operated hearts which reached statistical significance at 14 dpci (*p*=0.018) (Fig.1B).

### Neuropilins are upregulated in the epicardium and the endocardium following cryoinjury

We used *in situ* hybridization to delineate the spatio-temporal expression of neuropilins following cryoinjury (Fig. 1C). The specificity of the *nrp* RNA probes was initially analyzed in zebrafish embryos (Fig.S2), confirming expression patterns similar to previous observations in the literature (47, 48). In control sham-operated adult zebrafish hearts, *nrp1a* was expressed by the epicardium and *nrp2b* was widely expressed by the myocardium (Fig.1C). One day post cryoinjury, *in situ* hybridization revealed mRNA up-regulation of all neuropilin isoforms in the epicardium and at the interface between the healthy myocardium and the injured tissue. At 3 dpci, both *nrp1a* and *nrp2a* were strongly and more widely upregulated by the activated epicardium, while *nrp1b* was expressed at the injury border. At 14 dpci, strong expression of *nrp1a* persisted in the epicardium adjacent to the injured area and *nrp1b* was localized in the epicardium and the endocardium contiguous to the injured area. By 60 dpci, when heart regeneration was largely complete, expression of all *nrp* isoforms had returned to basal expression levels, which correlated with the gene expression data (data not shown).

To identify the Nrp1-expressing cells within the regenerating heart, we used co-immunofluorescent staining with specific endothelial, myocardial, and epicardial markers. In *tg(fli1a:EGFP)^y1^* zebrafish in which EGFP is specifically expressed in vascular endothelial cells, Nrp1 was co-expressed by *fli1a*-EGFP-expressing cells in sham hearts consistent with expression of Nrp1 by coronary vessels and endocardium (Fig.2A, upper row). Nrp1 expression was also evident in *fli1a*-EGFP-expressing neovasculature and activated endocardium at the injured area in cryoinjured hearts (Fig.2A, lower row). These observations were supported by immunostaining of *tg(kdrl:mCherry)^s896^* transgenic fish, in which mCherry expression is driven by the promoter for the endothelial VEGF receptor, *kdrl.* Nrp1 immunostaining at 7 dpci in *tg(kdrl:mCherry)^s896^* fish showed co-expression of mCherry-positive endocardium and Nrp1 (Fig.S3). Furthermore, neovascularization was observed as early as 1dpci in *tg(fli1a:EGFP)^y1^* fish, consistent with previous findings (25), and these early neovessels also demonstrated Nrp1 expression (Fig.S4). Nrp1 expression by tropomyosin-positive cardiomyocytes was low in control sham-operated hearts (Fig.2B, upper row). However, following cryoinjury, Nrp1 was expressed by a small population of cardiomyocytes located within the sub-epicardial layer at the lesion (Fig. 2B, lower row).

**Figure 2.**
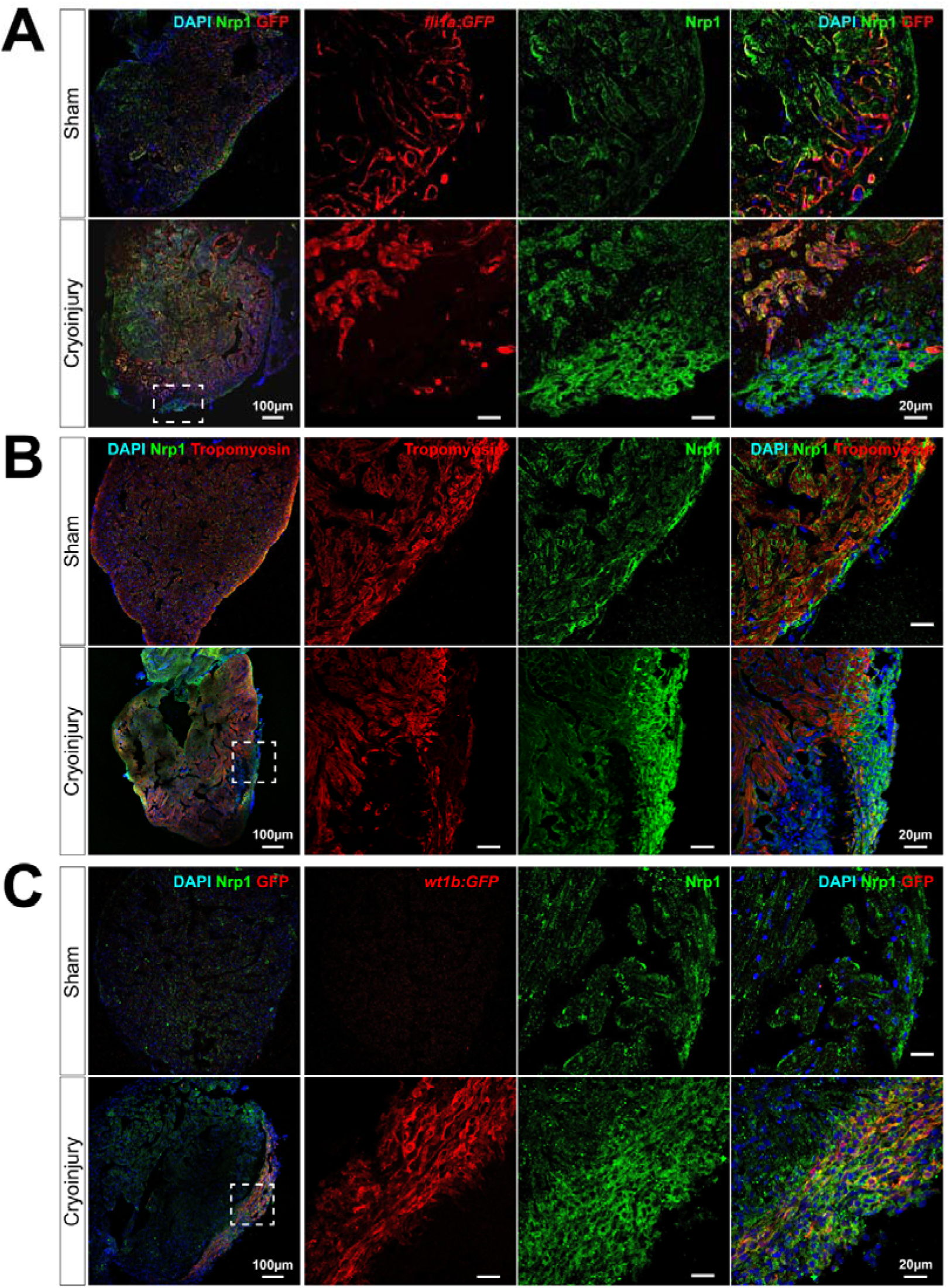
Nrp1 is expressed by the endocardium and the epicardium in sham and cyroinjured hearts. Immunostaining of 7 days post sham-operated (upper rows) and cryoinjured (lower rows) hearts. *Tg(fli1a:EGFP)^y1^* **(A**), Wild-Type fish immunostained for tropomyosin **(B**), and *tg(wt1b:EGFP)^li1^* **(C**) zebrafish hearts were used to identify endothelium, myocardium and activated epicardium, respectively. Overlay of the two colors are displayed with DAPI nuclei staining. White dotted boxes denote location of enlarged images. (n ≥ 3) V - ventricle, A -atrium, *ba* - *bulbus arteriosus*.

Epicardial expression of Nrp1 was examined in *tg(wt1b:EGFP)^li1^* zebrafish, in which EGFP expression is controlled by the promoter for the activated epicardial marker, Wilms’ tumor 1b (*wt1b*). In sham-operated control hearts of *tg(wt1b:EGFP)^li1^* zebrafish, no detectable expression of EGFP was observed (Fig.2C, upper row). In contrast, we observed high levels of colocalization between Nrp1 and EGFP in the epicardium covering the lesion in cryoinjured *tg(wt1b:EGFP)^li1^* zebrafish (Fig.2C, lower row). Furthermore, immunofluorescent staining of Wt1 and Nrp1 in cryoinjured Wild-Type zebrafish revealed a strong co-localization of Wt1 with Nrp1 in the epicardium adjacent to the injured area at 3dpci (Fig.S5).

### *Nrp1a* mutant zebrafish (*nrp1a^sa1485/sa1485^*) display delayed heart regeneration following cryoinjury

The marked upregulation of Nrp1 mRNA and protein at the borders of healthy and cryoinjured myocardium, and the expression of Nrps by the endocardium and the activated epicardium suggested a role for Nrps in heart regeneration, particularly in the activated epicardium. Because of the clear epicardial and endocardial expression of *nrp1a* after myocardial injury, we assessed the role of this isoform using the *nrp1a^sa1485/sa1485^* homozygous mutant zebrafish. This mutant carries a non-sense mutation (tyrosine to ochre, TAA) at amino acid (aa) 206 (full length, 916 aa) in the second CUB domain of the *nrp1a* gene, resulting in the generation of a non-functional and truncated soluble N-terminal fragment (Fig.3A). In these mutant fish, loss of *nrp1a* has been shown to induce axons to misproject to the dorsal and anterior dorsal zone protoglomerulus (49). *Nrp1a^sa1485/sa1485^* mutant fish were viable, born at expected Mendelian ratios (Fig.3B), displayed no obvious abnormal phenotype (Fig.S6A), and their body lengths and heart sizes were very similar to those of Wild-Type fish (Fig.S6B,C,D). In the *nrp1a^sa1485/sa1485^*fish, *nrp1a* expression was reduced at both the mRNA (Fig.3C and Fig.3D) and protein (Fig.3E) levels. Following cryoinjury, the extent of lesions in *nrp1a^sa1485/sa1485^* and Wild-Type fish hearts were similar, affecting 22.6 ± 5.2% (S.E.M) and 25.2 ± 5.5% (S.E.M) of the ventricle, respectively. AFOG staining was used to quantify the extent of the injury in both Wild-Type and *nrp1a^sa1485/sa1485^* fish over 60 days (Fig.4B). By 60 dpci, the injured area was almost cleared and new healthy myocardium had replaced the damaged tissue in Wild-Type fish (Fig.4A,B). A reduction in the extent of heart repair was observed from as early as 7 dpci in the *nrp1a^sa1485/sa1485^* hearts compared with Wild-Type fish (Fig.4A, B). Whereas fibrin deposits (red staining in injury area) were cleared from the injury scars in Wild-Type fish by 14 dpci, fibrin deposits were still evident at 30 and 60 dpci in the *nrp1a^sa1485/sa1485^* mutants (Fig.4A). Quantification of the size of the cryoinjuries using two-way ANOVA revealed a significant delay in the regeneration of mutant hearts in comparison to Wild-Type hearts (*p*=0.038). Furthermore, specific time-point comparison of injury sizes at 14 and 30 dpci showed that a significantly larger injury persisted in mutant hearts compared to Wild-Type hearts (*p*=0.0064 and 0.02, respectively) (Fig.4B). Differences between *nrp1a^sa1485/sa1485^* and Wild-Type hearts were also observed in the regeneration of the cortical layer and wound closure. In Wild-Type zebrafish hearts regeneration typically led to formation of a continuous layer of cardiomyocytes enclosing the residual collagen scar, resulting in complete wound closure in the advanced stages of regeneration (30 and 60 dpci). In contrast, a larger proportion of mutant hearts at 30 and 60 dpci retained open wounds without complete closure of the lesion (Fig.4C).

**Figure 3.**
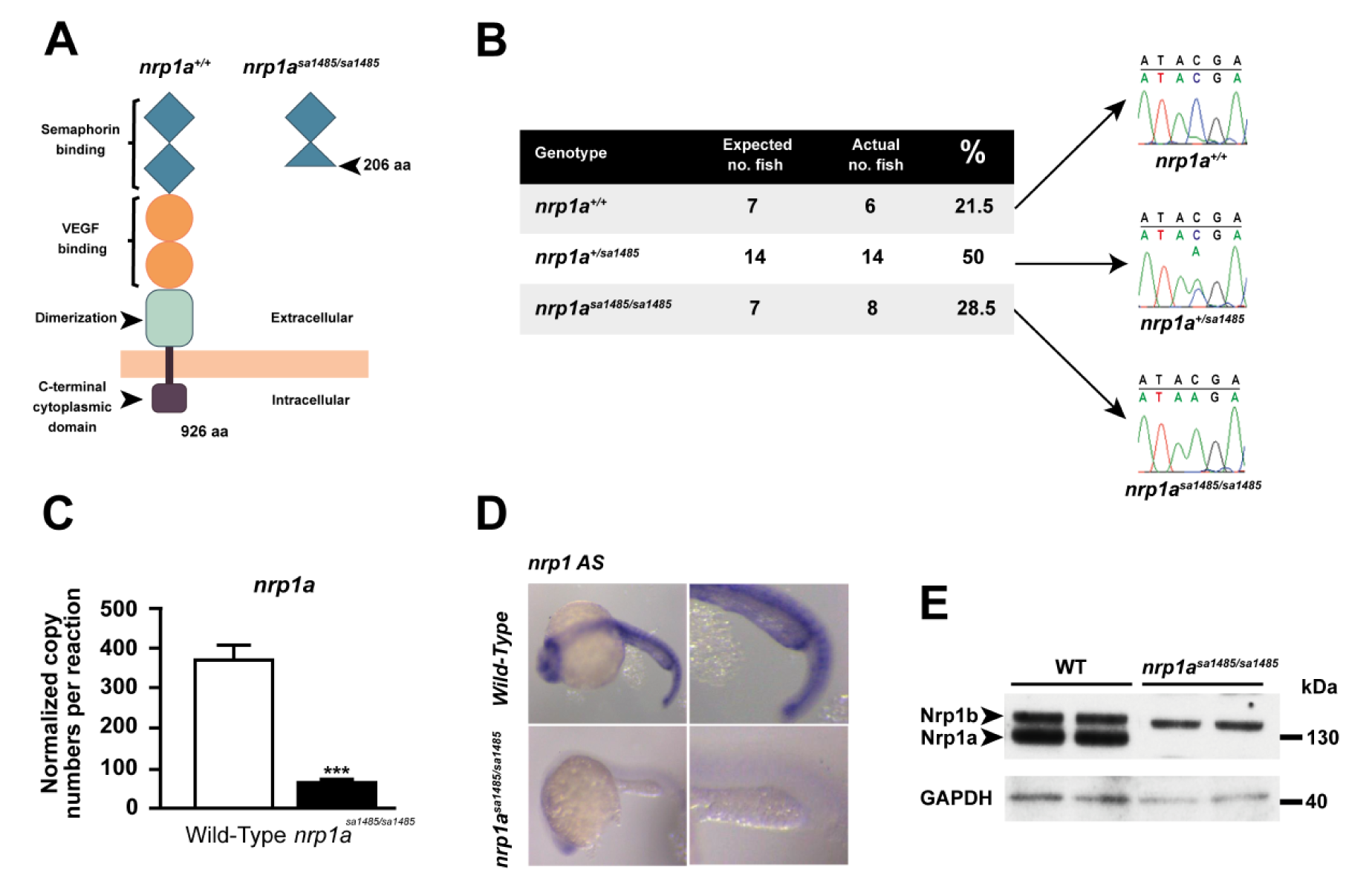
*nrp1a^sa1485/sa1485^* mutant fish characterization. **(A)** Diagram of Wild-Type (WT) (left) and *nrp1a^sa14854/sa1485^* mutant fish (right) Nrp1a structure. The point mutation results in the generation of a premature stop codon at amino acid 206 resulting in a truncated Nrp1a fragment. **(B)** Sequencing chromatograms of Wild-Type fish, heterozygous *nrp1a^sa1485/+^* and homozygous *nrp1a^sa1485/sa1485^* mutant fish. An early stop codon (nonsense mutation) TAA, replaces the Wild-Type TAC codon at amino acid 206. The genotypes of 14 zebrafish embryos 48 hours post fertilization (hpf) were compared against the expected Mendelian ratio after heterozygous fish incross. **(C)** Absolute RT-qPCR of Wild-Type (white bar) or *nrp1a^sa14854/sa1485^* homozygous mutant (black bar) uninjured adult zebrafish hearts under basal conditions. Results are represented as means of normalized copy numbers per reaction ± S.E.M (error bars) ^⋆⋆⋆^ p<0.005 *n* = 4 with each *n* being a pool of 3 ventricles. *nrp1a* expression is significantly decreased in *nrp1a^sa1485/sa1485^* samples. **(D)** *Nrp1a* anti-sense (AS) *in situ* hybridization of Wild-Type (upper row) or *nrp1a^sa1485/sa1485^* homozygous mutant (lower row) embryos 24 hpf. Black dotted boxes indicate magnified region. *Nrp1a* expression is clearly decreased in *nrp1a^sa14854/sa1485^* samples. **(E)** Western blot of Wild-Type (WT) or *nrp1a^sa1485/sa1485^* homozygous mutant uninjured adult zebrafish ventricle lysates. Lysates were immunoblotted for Nrp1 cytoplasmic domain and Gapdh. Note the absence of C-terminus detection of Nrp1a in the *nrp1a^sa1485/sa1485^* samples.

**Figure 4.**
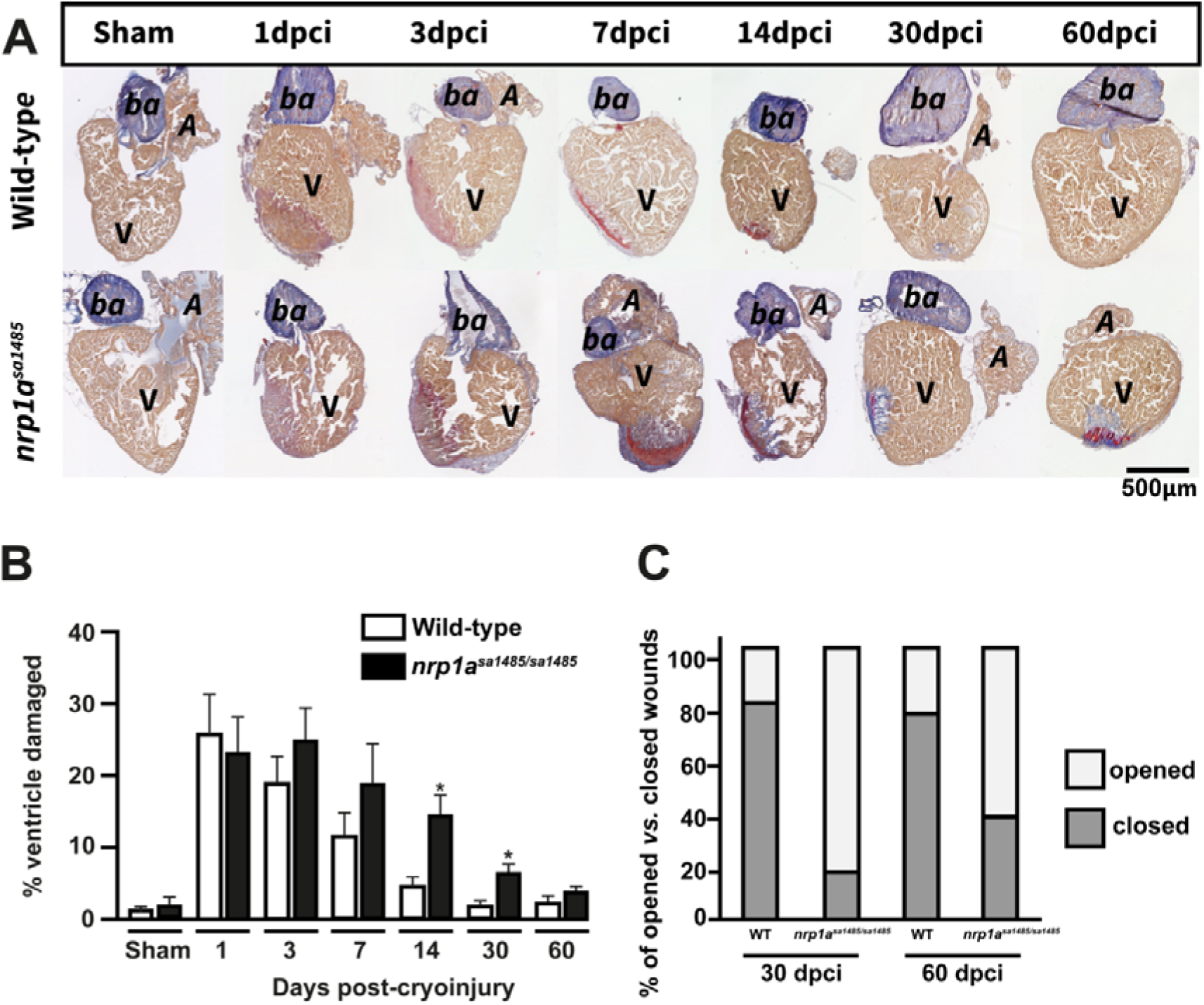
Recovery from cryoinjury is delayed in *nrp1a^sa1485/sa1485^* mutants. **(A)** Heart sections from Wild-Type and *nrp1a^sa1485/sa1485^* mutant fish obtained at 1,3, 7, 14, 30 and 60 dpci and stained with AFOG to identify the injured region **(B)** Cryoinjured areas were measured and represented as mean percentage of total ventricle area ± S.E.M (n = 4-8) ^⋆^*p* ≤ 0.05. **(C)** The compact myocardium recovers following cryoinjury and encases the scar tissue, this is defined as a closed wound; whereas a scar exposed to the surface is defined as an open wound. Wound closure was examined in Wild-Type and *nrp1a^sa1485/sa1485^* mutant hearts at 30 and 60 dpci and open *vs* closed wounds were counted as percentage of total number of hearts (n=4-8), *A* – atrium, *ba* – *bulbus arteriosus*, V – ventricle.

### Revascularization of the cryoinjured heart tissue is impaired in *nrp1a* mutant zebrafish (*nrp1a^sa1485/sa1485^*)

The impact of loss of functional Nrp1 on revascularization in cryoinjured hearts was examined by generating *nrp1a^sa1485/sa1485^* mutants *in tg(fli1a:EGFP)^y1^* zebrafish in which endothelial-specific GFP expression is driven by the *fli1a* promoter. Angiogenesis occurs rapidly following heart injury in zebrafish, with a marked neovascular response evident as early as 1 dpci (25). Therefore, we compared the extent of angiogenesis in control *tg(fli1a:EGFP)^y1^* and compound *tg(fli1a:EGFP)^y1^:nrp1a^sa1485/sa1485^* mutant zebrafish at 1 and 3 dpci. GFP-positive vessels were clearly identified within the injured area at 1 and 3 dpci in both Wild-Type and mutant zebrafish (Fig.5A and B). However, the sa1485 mutation was associated with a significant, 3 to 4-fold reduction in the extent of neovascularization. At 1 dpci, the average number of coronary vessels found within each microscopic field (32,625μm^2^) of the injury was 13 in Wild-Type zebrafish compared to 3 in *nrp1a^sa1485/sa1485^* mutant zebrafish (p=0.0087) (Fig.5A). At 3 dpci, the average number of newly formed vessels within the injured area per microscopic field of the injury was 19 in Wild-Type zebrafish compared to 6 in *nrp1a^sa1485/sa1485^* mutant zebrafish (p=0.0258) (Fig.5B).

**Figure 5.**
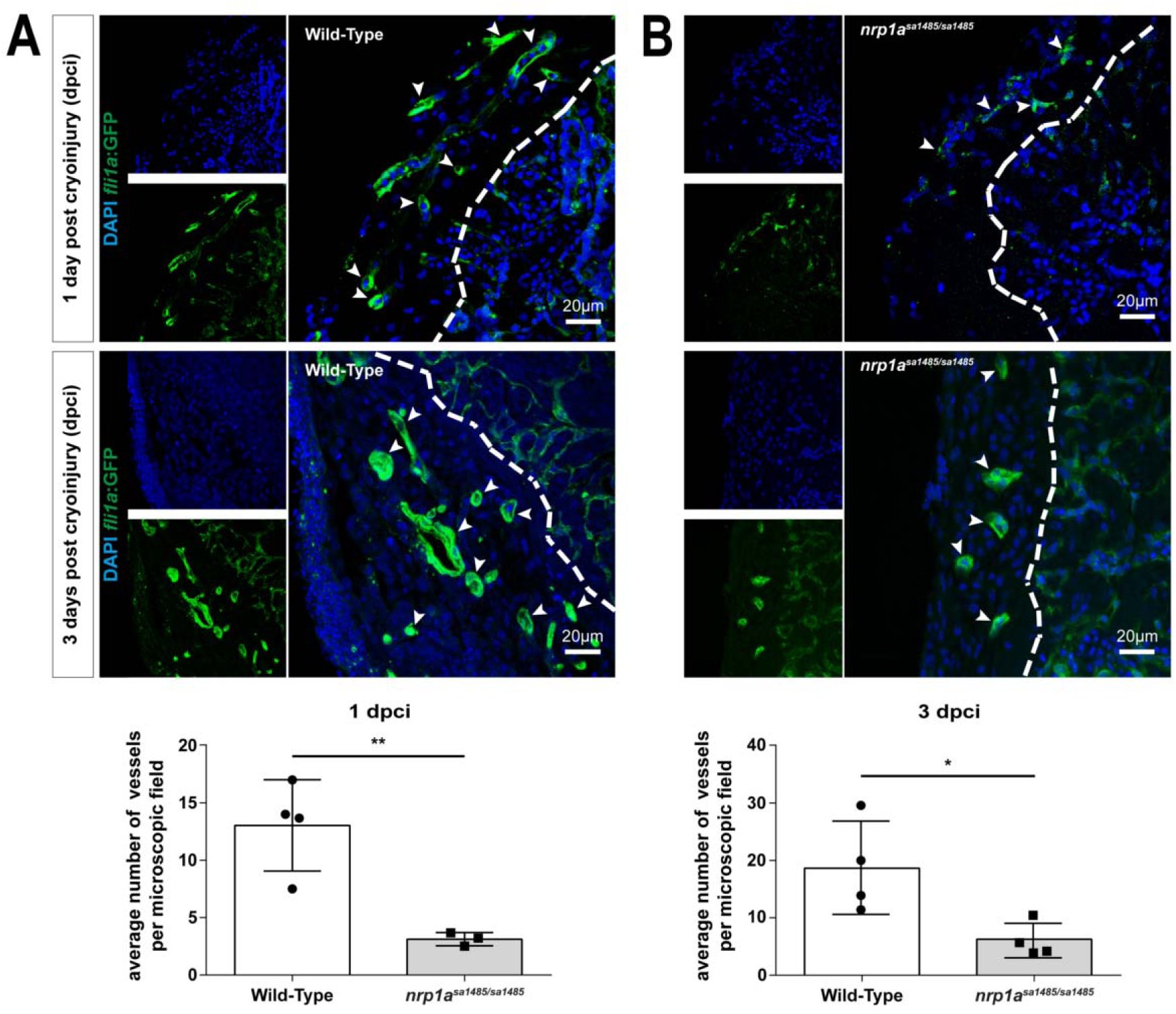
Neovascularization of the cryoinjured area is impaired in *nrp1a^sa1485/sa1485^* mutants. Blood vessels in either Wild-type **(A)** or *nrp1a^sa1485/sa1485^* **(B)** *tg(fli1a:EGFP)^y1^* zebrafish at 1 day (upper row) and 3 days (lower row) post cryoinjury were identified in heart sections by GFP immunofluorescence in vascular structures. Heart sections were also counterstained with DAPI. The dashed white line delineates the border of the area of injury. White arrows indicate blood vessels. GFP-positive vessels were then quantified: ^⋆⋆^ p < 0.01, ^⋆^ p < 0.05 for Wild-Type *versus nrp1a^sa1485/sa1485^* hearts; individual data points represent vessel counts in individual hearts.

We further examined angiogenesis in cryoinjured hearts from Wild-Type and *nrp1a^sa1485/sa1485^* mutant zebrafish at 3 dpci by immunofluorescent staining for the endothelial-specific marker Tie2. Use of Tie2 immunostaining as a reliable method to identify neovessels was verified by comparing the number of vessels within the cryoinjury of *tg(fli1a:EGFP)^y1^* zebrafish identified either using Tie2 or GFP immunostaining (data not shown). The sa1485 mutation was associated with a significant reduction in the extent of neovascularization (Fig.S7). At 3 dpci, the average number of Tie2-positive vessels was 4.776 per field (n=7) in Wild-Type zebrafish compared to 2.557 in *nrp1a^sa1485/sa1485^* mutant zebrafish (n=6) (p=0.0058) (Fig.S7).

### Epicardial activation is inhibited in *nrp1a^sa1485/sa1485^* heart

We next addressed whether the delayed heart regeneration caused by loss of functional Nrp1 in *nrp1a^sa1485/sa1485^* zebrafish could be due to an impact on activation of the epicardium and subsequent epicardial regeneration. Consistent with this possibility, our data showed epicardial upregulation of Nrp1 adjacent to the injured area indicated by strong co-localization of Nrp1 with Wt1b, a specific marker for epicardium activation, at 3 dpci (see Fig.2, Fig.S8A), a time coincident with occurrence of robust epicardial activation during zebrafish heart regeneration (50). To investigate this hypothesis, we examined epicardial activation in cryoinjured hearts of Wild-Type and *nrp1a^sa1485/sa1485^* mutant *tg(wt1b:EGFP)^li1^* zebrafish. Analysis of hearts at 3 dpci revealed a strong decrease in GFP expression under the control of the *wt1b* promoter in the *nrp1a^sa1485/sa1485^* mutants as compared to Wild-Type (Fig.6A). Quantification of GFP-positive cells within the epicardium covering the cryoinjured area confirmed a marked and significant reduction in the number of GFP-expressing activated epicardial cells in *nrp1a^sa1485/sa1485^* mutants (14.08 *versus* 26.4 for *nrp1a^sa1485/sa1485^* and Wild-Type, respectively; *p*=0.0071)(Fig.6B). We also investigated whether loss of functional Nrp1 impaired the proliferation of activated epicardial cells, using PCNA staining. *Nrp1a^sa1485/sa1485^* hearts showed no statistically significant reduction in the numbers of proliferating cells compared to Wild-Type hearts (76.88% versus 91.46% for *nrp1a^sa1485/sa1485^* and Wild-Type, respectively; *p*=0.2138)(Fig.6C and D).

**Figure 6.**
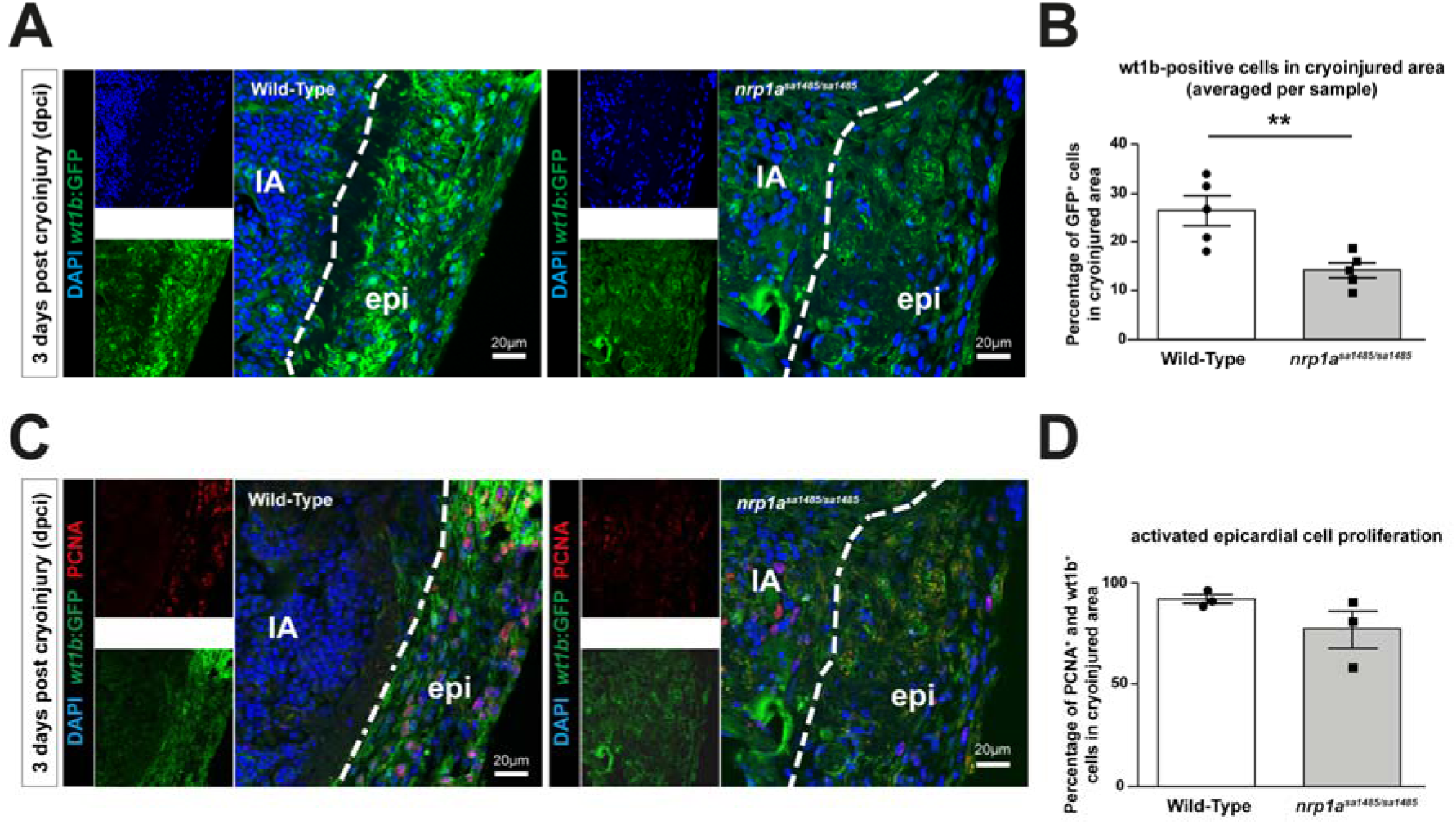
Epicardial activation is decreased in *nrp1a^sa1485/sa1485^* hearts following cryoinjury. Wild-Type and *nrp1a^sa1485/sa1485^* mutant cryoinjured fish on the *tg(wt1b:EGFP)^1i1^* background were analyzed for epicardial activation at 3 dpci by identification of *wt1b*:EGFP-positive cells **(A,C)**, and were also stained with DAPI **(A,C)**, and anti-PCNA antibody **(C)**, as indicated. The percentages of *wt1b*:EGFP-positive cells in the area of cryoinjury (indicated by the dashed line) were quantified in Wild-Type and *nrp1a^sa1485/sa148S^* mutant fish **(B)**: ^⋆⋆^ p < 0.01, ^⋆^ p < 0.05 for WT *vs nrp1a^sa1485/sa1485^* hearts; individual data points represent percentages in individual hearts. The percentages of cells positive for *wt1b*:EGFP and PCNA were also quantified in the area of cryoinjury **(D)**: individual data points represent percentages in individual hearts. (IA – injury area, epi – epicardium white dotted line delineates injury/epicardial border).

Additionally, we examined whether cardiomyocyte proliferation was affected in *nrp1a^sa1485/sa1485^* mutant hearts by determining the number of tropomyosin-positive cells that were also PCNA-positive. The results showed a modest reduction in tropomyosin+/PCNA+ cells in *nrp1a^sa1485/sa1485^* mutants as compared with Wild-type hearts (26.65% versus 36.7%; p=0.2366, respectively)(Fig.S9) but this was not statistically significant.

### Epicardial expansion and activation of cryoinjury-induced *nrp1a^sa1485/sa1485^* heart explants is impaired

We next examined the role of Nrp1 in epicardial activation in an *ex vivo* heart explant model. Immunofluorescent staining of Nrp1 in explants of *tg(wt1b:GFP)^li1^* ventricular apexes collected at 5 dpci and cultured *in vitro* for 7 days showed perinuclear, cytoplasmic and membrane localization (Fig.S8B). *Wt1b:EGFP* expression was variable in these explants and was strongly expressed by a subpopulation of explanted epicardial cells (top row, Fig.S8B).

As shown in Fig.S10A and C, *wt1b*:EGFP expression was increased in explant outgrowths from cryoinjured as compared with those from control, sham-operated hearts and, similar to resected hearts (51), explants from Wild-Type cryoinjured hearts generated greater outgrowth in comparison with those from Wild-Type sham-operated hearts (Fig.7A,B and Fig.S10A,B). We next assessed the role of Nrp1a in injury-induced epicardial activation using epicardial explants from Wild-Type and *nrp1a^sa1485/sa1485^ tg(wt1b:EGFP)^li1^* zebrafish (51). Epicardial expansion from cryoinjury-induced *nrp1a* heart explants was markedly impaired compared with Wild-Type explants (Fig.7A,B) and resembled that of sham-operated hearts, confirming our observations of reduced GFP-positive epicardial cells in sections from cryoinjured *nrp1a^sa1485/sa1485^ tg(wt1b:EGFP)^li1^* zebrafish hearts (Fig. 6A,B). Furthermore, *nrp1a^sa1485/sa1485^* mutants showed a marked decrease of GFP expression in comparison with the Wild-Type fish, both at the edge of the explant and within the region closest to the heart (Fig.7C), providing further support for a loss of epicardial activation in the *nrp1a^sa1485/sa1485^* mutants in comparison to Wild-Type hearts.

**Figure 7.**
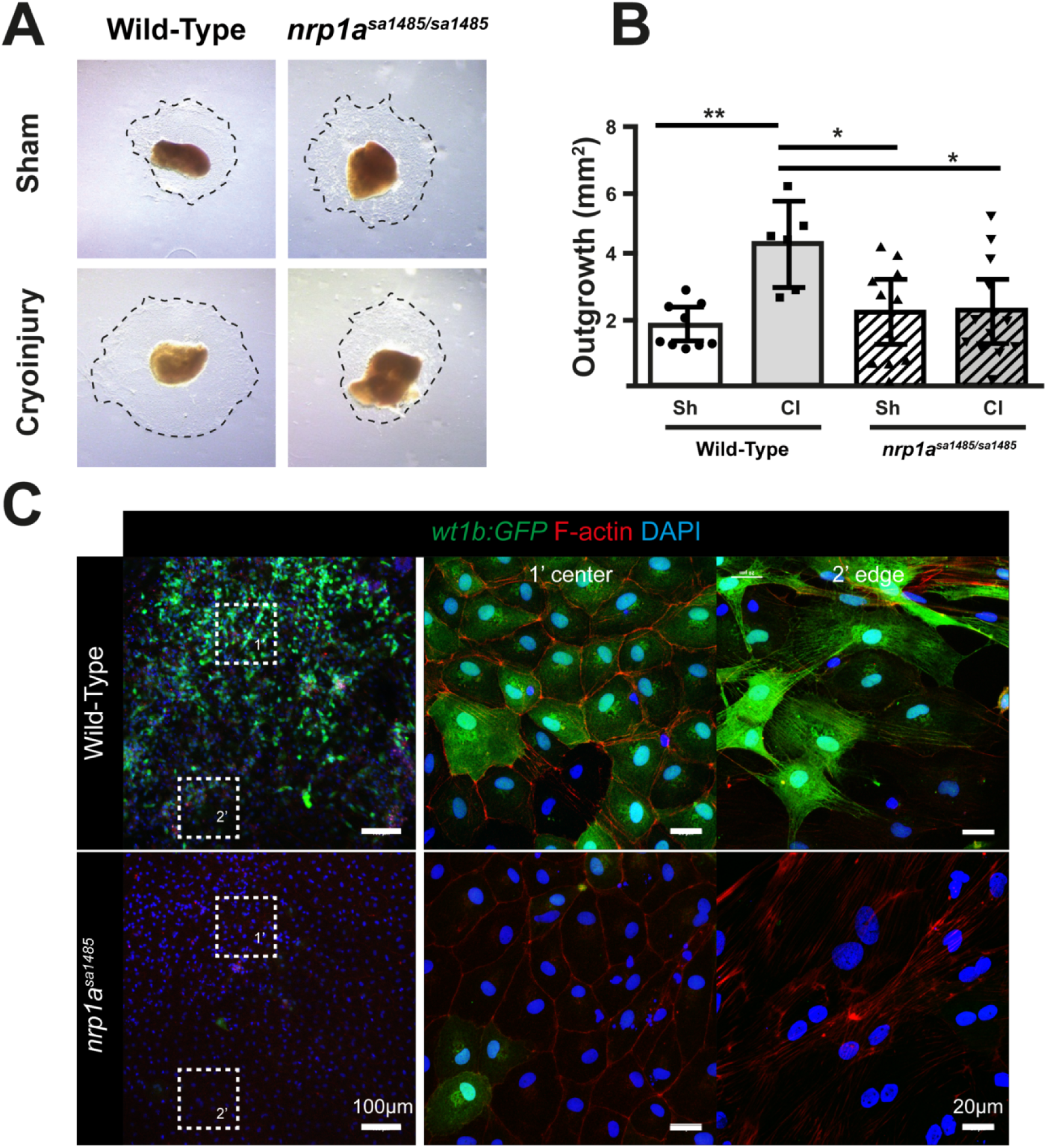
Epicardial cryoinjury induced expansion and activation are impaired in *nrp1a^sa1485/sa1485^* mutants. The apices of Wild-Type and *nrp1a^sa1485/sa1485^* zebrafish ventricles were collected 5 days post sham surgery or cryoinjury and cultured on fibrin gels for 7 days. **(A)** Epicardial cells migrate into the fibrin gels (dotted black lines). **(B)** Epicardial outgrowths were measured for each condition after 7 days culture, data is represented as mean outgrowth area (mm^2^) ± S.E.M (n = 6-12) ^⋆^*p*<0.05, ^⋆⋆^*p*<0.01, scale bar 1 mm. **(C)** Epicardial explant recovered from Wild-Type and *nrp1a^sa1485/sa1485^ tg(wt1b:GFP)^111^* cryoinjured fish at 5 dpci were left to grow on fibrin gels for 7 days and stained for GFP. GFP fluorescence was observed at 100x **(left column)** and 400x magnification at the center and the periphery of the explants (**middle** and **right columns** respectively).

## Discussion

Epicardial activation and angiogenesis are processes essential for zebrafish heart regeneration following injury. During these processes, revascularization and injury-induced epicardial-to-mesenchymal transition (EMT) are driven by Vegf, Fgf, and Pdgf (10, 17, 18), which are all ligands for the cell surface receptor Nrp1. The role of Nrps has not previously been characterized in zebrafish heart regeneration, though NRPs are essential for angiogenesis and increasingly implicated in EMT in other contexts in mammalian species (26, 31, 33, 42, 43). Here, we demonstrate for the first time that *nrp1* and *nrp2* are upregulated in response to cardiac injury and that *nrp1a* plays an essential role in both revascularization and epicardial activation and migration, processes that are essential for the regeneration of the zebrafish heart.

*Nrp1a*, *nrp1b*, *nrp2a*, and *nrp2b* mRNAs were all strongly upregulated in the zebrafish heart 1-3 days after cryoinjury, coinciding with the time of epicardial activation, which occurs very early following cardiac injury (16, 17). Increased protein expression of both Nrp1 isoforms also occurred 3-14 days following cryoinjury, further supporting the conclusion that Nrp1 is upregulated in the early regenerating heart. Our results also revealed a striking spatio-temporal upregulation of the *nrp* isoforms. Specifically, *nrp2a* was strongly upregulated in the epicardium and endocardium proximal to the injury at 1 and 3 dpci, whereas *nrp1a* was strongly upregulated in the same regions at 1,3 and 14dpci. These findings support a sustained role for these isoforms in heart regeneration, particularly in the epicardial activation phase which takes place during the first 3-7 days of regeneration. At present, the lack of suitable reagents made further detailed studies of Nrp2 problematic. However, a role for Nrp1 in epicardial activation in response to heart injury was further supported by immunofluorescent staining demonstrating co-localization of Nrp1 with both endogenous Wt1 and with GFP under the control of the *wt1b* promoter, an embryonic gene that is upregulated following cardiac injury and is an activated epicardium marker (23, 52).

Analysis of mutant *nrp1a^sa1485/sa1485^* fish lacking expression of full-length Nrp1a provided direct evidence that Nrp1a is required for zebrafish heart regeneration. *Nrp1a^sa1485/sa1485^* fish displayed no morphological or pathological phenotype. It was previously reported that knockdown of *nrp1* using morpholino oligomers produces a lethal phenotype in zebrafish embryos (44). The absence of embryonic lethality in *nrp1a^sa1485/sa1485^* fish as compared with *nrp1a* morpholino knock-down probably reflects redundancy due to adaptive mechanisms relying on compensation by *nrp1b* and may also suggest morpholino off-target effects in *nrp1a* morphants (53). Following cardiac damage, *nrp1a^sa1485/sa1485^* mutants exhibited a significantly reduced regenerative response in comparison with Wild-Type controls. The importance of *nrp1a* for heart regeneration was demonstrated by the delayed and incomplete removal of fibrin deposits essential for the scar resolution process in *nrp1a^sa1485/sa1485^* mutant fish. Given that myocardial proliferation was not significantly affected in *nrp1a^sa1485/sa1485^* mutant fish (Fig.S9), delayed wound closure in *nrp1a^sa1485/sa1485^* fish indicates a failure of the myocardium to migrate efficiently towards the subepicardial layer after cryoinjury. Together, these findings provide strong support for the conclusion that *nrp1a* is required for zebrafish heart regeneration following cryoinjury. Since we examined the loss of only the *nrp1a* isoform, due to anticipated embryonic lethality of a double *nrp1a* and *nrp1b* knock out, it is possible that Nrp1 loss may have an even more prominent role in heart regeneration in species that did not undergo genome duplication. Furthermore, our data indicated prominent epicardial expression of *nrp2*, indicating a possible role of Nrp2 isoforms in epicardial activation and heart regeneration, something that warrants further investigation.

Activation of the epicardium and subsequent regeneration involves multiple cellular processes, including cell migration, proliferation, and EMT. It is well established that NRP1 modulates cell migration in diverse mammalian cell types (29, 54-57). The conclusion that *nrp1a* is important for zebrafish epicardial migration is supported by our finding that *ex vivo* outgrowth from epicardial explants of *nrp1a^sa1485/sa1485^* hearts was also impaired. In contrast, we observed no effect on epicardial cell proliferation in cryoinjured *nrp1a^sa1485/sa1485^* hearts, indicating that *nrp1a* is not critical for proliferation, in line with studies of NRP1 function in mammalian cells (29). However, we cannot preclude the possibility that one of the other *nrp* isoforms has a role in epicardial proliferation.

Reduced *in vitro* expansion of *nrp1a*-deficient epicardial cells is likely due to impaired detection of cellular cues promoting migration. Consistent with this possibility, we observed upregulation of *nrp* isoform expression concomitant with increased expression of *tgfb*, *pdgfab*, *vegfc*, and the receptor *pdgfrb*, chemotactic factors and receptors implicated in zebrafish heart regeneration, and also shown to act as ligands and co-receptors for NRP1 in mediating mammalian cellular functions. Interestingly, using the *(wt1b:EGFP)^li1^* transgenic fish line, we also noted that the *nrp1a^sa1485/sa1485^* epicardial cells failed to re-express the *wt1b* embryonic marker *in vitro* as well as *in vivo.* Previously, Gonzalez-Rosa *et al.* (58) demonstrated the importance of the wt1b:EGFP+ epicardial derived cells (EPDCs) in the regeneration process which gave rise to perivascular fibroblasts and myofibroblasts and also participated in the regeneration process by secreting essential pro-angiogenic paracrine factors (58). The decreased number of GFP-expressing (and therefore *wt1b*^+^) EPDCs by the *nrp1a^sa1485/sa1485^* hearts explains their delayed regeneration in comparison to WT hearts.

Our study also revealed Nrp1 upregulation following cardiac damage by the activated endocardium, which undergoes endothelial to mesenchymal transition (endoMT) in response to injury (16), by the neovasculature, and by some subepicardial cardiomyocytes known to be a primary source of new myocardium (59). Following injury, these cells acquire a migratory phenotype to contribute to the regenerative processes in the heart.

Nrp1 has an essential role in angiogenesis in mammalian and zebrafish development, and is also required in post-natal and adult angiogenic processes (33, 41). Marin-Juez *et al.* recently reported transient upregulation of *vegfaa* at 1 dpci, with a return to baseline expression by 3 dpci, and showed an important role for *vegfaa* in inducing rapid early revascularization of the injured heart (25). Our data showed a trend towards increased *vegfaa* expression at 1 dpci, using RT-qPCR, although this was not statistically significant, unlike the concomitant changes in *nrp1.* Similarly to its major endothelial ligand *vegfaa*, the main Nrp1 co-receptor *kdrl* was also not significantly upregulated. However, we observed revascularization of the injured area as early as 1dpci, in line with previous findings (25). These neovessels also expressed Nrp1 and studies in *nrp1a^sa1485/sa1485^* mutants co-expressing *fli1a:EGFP^y1^* demonstrated a role for *nrp1a* in the revascularization of the cryoinjured area. As expected, the loss of Nrp1a reduced the number of neovessels in the regenerating heart of *nrp1a^sa1485/sa1485^* fish in comparison to their Wild-Type counterparts. While our findings are consistent with a role for Nrp1 in mediating Vegfaa-driven angiogenesis in the regenerating heart, recent findings indicate that the role of NRP1 in developmental angiogenesis may be largely independent of VEGF, since NRP1 mutations which prevent VEGF-A binding impair post-natal angiogenesis but are compatible with normal embryonic development (60). It is therefore plausible that the angiogenic role of Nrp1 in revascularization of the regenerating zebrafish heart is also mediated via binding of other ligands to Nrp1 (26-28).

This study establishes a novel role for Nrp1 in epicardial activation and angiogenesis during zebrafish heart regeneration following injury. Further work to elucidate the extracellular ligands for Nrp1 in epicardial and endothelial cells and the signaling pathways that mediate its role further downstream will shed new light on the mechanisms involved in epicardial activation in heart regeneration.

## Materials and Methods

An extended version of the Material and Methods section is available as supplementary information online.

### Zebrafish husbandry and cryoinjury

Procedures were performed in accordance with the Animals (Scientific Procedures) Act 1986, and husbandry was regulated by the Central University College London fish facility. Cryoinjury procedure was carried out as described in (61), and more details are provided in the supplementary ‘Materials and Methods’.

### RT-qPCR

Ventricles from corresponding time-points and treatments were pooled for RNA extraction using the RNeasy Mini Kit (Qiagen). RNA was reverse transcribed using the QuantiTect^®^ Reverse Transcription Kit (Qiagen). All primers (details are described in table S1) and standards were purchased from qStandard©.

### Histological procedures

*In situ* hybridization, immunofluorescence and Acid Fuchsin Orange G (AFOG) procedures are extensively described in the supplementary information file.

### Fibrin gel heart explants

*In vitro* epicardial cell outgrowth experiments were performed as previously described (62). The apex of cryoinjured and sham-operated zebrafish hearts were isolated 5 days post-surgery and placed firmly on set fibrin gel matrices, ensuring epicardial surface contact with the gel. Medium was changed every 2 days and cells were cultured for 7 days before harvesting epicardial outgrowths for protein extraction or immunofluorescence imaging.

### Statistical analysis

Results are presented as mean ± standard error of the mean (S.E.M). Experimental repeat *n* values are indicated as individual data points in graphs or specified in figure legends. All data were analyzed using Graphpad prism 6.0. qPCR and AFOG cryoinjury area data were analyzed for statistical significance using two-way ANOVA. Student’s t-test analysis was applied to all other data sets. Statistical significance values are indicated in figure legends whereby data were considered significant for *p*<0.05. Immunostaining data was analyzed by a blinded investigator.

## Acknowledgements

We thank Juan Manuel González Rosa for teaching us the cryoinjury model, Dr. Gaia Gestri for her help with the ISH protocol and for providing us with the *tg(kdrl:mCherry)^s896^* fish, Dr. Shanie Budhram-Mahadeo for her advice to generate the ISH probes, Dr. Stefan Schulte-Merker for the pCRII vector used for generating the nrp2b ISH probe. We also would like to acknowledge Roisin Brid Doohan and Dr. Mathilda Mommersteeg for advising on histological procedures and Dr. Patricia De Winter and Dr. David Sugden (qStandard©) for qPCR services, University College London central fish facility for *in vitro* fertilization of *nrp1a^sa1485^* fish line and maintenance of fish stocks.

*Author contributions*
V.L and C.P-M. designed the study, performed experiments, analyzed and interpreted the data, and wrote the paper. I.Z. obtained funding, designed the study, interpreted the data, and wrote the paper. L.W. performed experiments, analyzed the data and wrote the paper, and J.S. performed experiments and analyzed the data. PF helped writing the paper and N.M-H assisted in the set-up of the cryoinjury model and provided reagents.

